# Potent immunogenicity and protective efficacy of a multi-pathogen vaccination targeting Zaire ebolavirus, Sudan ebolavirus, Marburg and Lassa viruses

**DOI:** 10.1101/2023.11.03.565549

**Authors:** Amy Flaxman, Sarah Sebastian, Sofia Appelberg, Kuan M Cha, Marta Ulaszewska, Jyothi Purushotham, Ciaran Gilbride, Hannah Sharpe, Alexandra J Spencer, Sagida Bibi, Daniel Wright, Stuart Dowall, Linda Easterbrook, Stephen Findlay-Wilson, Sarah Gilbert, Ali Mirazimi, Teresa Lambe

**Author notes:** These authors contributed equally to this work (Sarah Sebastian, Sofia Appelberg).

## Abstract

Viral haemorrhagic fevers (VHF) pose a significant threat to human health. In recent years, VHF outbreaks caused by Ebola, Marburg and Lassa viruses have caused substantial morbidity and mortality in West and Central Africa. In 2022, an Ebola disease outbreak in Uganda caused by Sudan ebolavirus resulted in 164 cases with 55 deaths. In February 2023, a Marburg disease outbreak was confirmed in Equatorial Guinea resulting in 15 confirmed and 23 suspected cases to date, with a second outbreak occurring concurrently in Tanzania. There are no clearly defined correlates of protection against these VHF, impeding targeted subunit vaccine development. Any vaccine developed should therefore induce strong and preferably long-lasting humoral and cellular immunity against these viruses. Ideally this immunity should also cross-protect against viral variants, which are known to circulate in animal reservoirs and cause human disease. We have utilized two viral vectored vaccine platforms, an adenovirus (ChAdOx1) and Modified Vaccinia Ankara (MVA), to develop a multi-pathogen vaccine regime against three filoviruses (Zaire ebolavirus, Sudan ebolavirus, Marburg) and an arenavirus (Lassa). These platform technologies have consistently demonstrated the capability to induce robust cellular and humoral antigen-specific immunity in humans, most recently in the rollout of the licensed ChAdOx1-nCoV19 /AZD1222. Here, we show that our multi-pathogen vaccines elicit strong cellular and humoral immunity, induce a diverse range of chemokines and cytokines, and most importantly, confers protection after lethal Zaire ebolavirus, Sudan ebolavirus and Marburg virus challenges in a small animal model.

**Author summary:** Outbreaks caused by Ebola and Lassa viruses have made headlines worldwide in recent years. Most recently, in 2023 a Marburg virus outbreak has claimed tens of lives with a high case fatality rate. As yet, no licensed vaccine exists to protect against this and other viral haemorrhagic fevers. An ideal vaccine would induce long-lasting immunity to, and protection from, viruses causing viral haemorrhagic fevers. We developed vaccines which can target multiple strains of Ebolavirus, the closely related Marburg virus and Lassa virus. The geographical ranges of these viruses overlap in West and Central Africa. We used viral vector platform technologies to generate these vaccines; ChAdOx1 has now been administered worldwide as part of COVID-19 vaccine rollouts, and MVA has been used in numerous clinical trials thus far. We found that both long lasting, antigen specific T cell and antibody responses were induced after vaccination. Lastly, we demonstrated these vaccines could protect small animals against challenge with Zaire ebolavirus, Sudan ebolavirus and Marburg virus.

## Introduction

Lassa fever (LF) is a viral haemorrhagic fever (VHF) whose aetiologic agent, Lassa virus (LASV), is a member of the Arenaviridae family. Each year, LASV is estimated to infect 100,000–300,000 humans with an estimated 5,000 deaths [1]. It is transmitted via contact with rodents and is endemic throughout most of West Africa. This geographical area overlaps with filovirus outbreaks, including the Zaire ebolavirus species (EBOV) and Sudan ebolavirus (SUDV) species, and the more recently discovered fruit bat reservoir of Marburg virus (MARV) [2]. Whilst VHFs caused by Ebolavirus and MARV are associated with high levels of morbidity and mortality, LF has a much lower-case fatality rate but causes a significant burden of disease in West Africa. Another similarity is the lack of available therapeutics or vaccines[3]. The need to develop therapeutic and prophylactic treatments against re-emerging and outbreak pathogens became starkly apparent during both the 2013–2016 West Africa EBOV epidemic[4] [5] and the 2020-2022 COVID-19 pandemic. Indeed, in recent years there have been numerous further outbreaks, including very large LF outbreaks in Nigeria in 2018 and 2019, with high case fatality rates (approximately 25%), the second largest outbreak of Ebola virus disease (EVD) in the Democratic Republic of the Congo in 2020, a SUDV outbreak in Uganda in 2022 and a Marburgvirus outbreak in Equatorial Guinea in 2023. All of these further underpin the critical need for preventive measures against VHF [6] [7].

Antibodies against viral surface proteins can prevent viral attachment and subsequent infection while a cellular immune response to the surface protein (as well as to internal viral proteins) can reduce disease severity and transmission in a number of infectious disease settings. As a result, viral surface proteins are the main antigenic targets in licensed vaccines against viral diseases today. The recombinant, replication competent vesicular stomatitis virus-based candidate vaccine (rVSV-ZEBOV), licensed in 2019, encodes the single EBOV surface glycoprotein (GP). This vaccine offers substantial protection against disease, shown in trials during the 2013-2016 and 2018 outbreaks[8]. In addition, a VSV-based LF vaccine candidate encoding the glycoprotein of LASV demonstrated protective efficacy in a non-human primate challenge against both a homologous and heterologous virus[9, 10], suggesting that LASV GP antigen is a tenable vaccine target. However, there are a limited number of vaccine platforms that can concurrently induce strong humoral and cellular immunity against viral antigens, while also having the cargo capacity to encode more than one antigen. By harbouring a cargo capacity of 7kb and inducing robust humoral and cellular antigen-specific immune responses after single administration [11], adenoviral vectors fulfil these requirements.

A multivalent vaccine approach, which is the WHO’s preferred target product profile, will ideally prevent disease caused by multiple filoviruses and other haemorrhagic fever viruses. Working towards this goal, we sought to develop multi-pathogen vaccines that can provide effective and long-lived protection against VHF caused by filoviruses and LASV. We demonstrate that our multi-pathogen adenoviral vectored vaccine induces a strong cellular and humoral immune response, which confers protection in both a lethal EBOV challenge in guinea pigs and in lethal EBOV, SUDV and MARV challenges in a type-I interferon receptor (IFNAR) knockout mouse model. However, it did not confer complete protection in a LASV challenge. Importantly, no evidence of immune competition between vaccine-encoded antigens is found after administration of our adenoviral-vectored multi-pathogen vaccine. The adaptive immune response after adenovirus immunisation can be further enhanced by a boost vaccination with a multi-pathogen Modified Vaccinia Ankara (MVA) vaccine, resulting in a strong adaptive immune response and a diverse chemokine and cytokine profile. These vaccines could be rapidly deployed to a target population when a viral haemorrhagic fever outbreak occurs.

## Methods

### Ethics statement

Mouse immunogenicity studies carried out under UK Home Office Project licences 30/2889 and P9804B4F1, were approved by the University of Oxford Animal Care and Ethical Review Committee and were carried out at the University of Oxford, Old Road Campus. Guinea pig procedures carried out under UK Home Office Project licence P82D9CB4B, were approved by the Public Health England (PHE) (now UKHSA) Animal Welfare Ethics Review Board (AWERB) and were carried out at PHE (now UKHSA), Porton Down, Salisbury. Animal work performed in the UK was carried out in accordance with the UK Home Office Animal Testing and Research Guidance as per the Animals (Scientific Procedures) Act 1986.

Mouse challenge studies were carried out under the Public Health Agency of Sweden’s, ethical licence 16676-2020, which were approved by Stockholm ethical committee for animal research. The study was carried out at the Karolinska Institute and at the Public Health Agency of Sweden. The mice were housed according to the Karolinska Institutes rules regarding animal care and observed at least once a day.

### Antigens

Amino acid sequences for antigens used in vector construction were as follows: EBOV (Zaire ebolavirus glycoprotein, Makona-Kissidougou-C15 GenBank: KJ660346.1), SUDV (Sudan ebolavirus glycoprotein, UniProtKB Q66814.1), MARV (Marburg virus glycoprotein, UniProtKB Q1PD50.1), LASV (Lassa virus glycoprotein (GPC), NCBI Reference Sequence: NP_694870.1). Antigen sequences were codon-optimised for *H. sapiens* and synthesised by Geneart (ThermoFisher, Germany).

### Vector construction

The derivation of the ChAdOx1 vector has been described before [12]. To generate recombinant vectors, antigen cassettes consisting of a TetR-repressible CMV promoter, antigen coding sequence, and polyA sequence were inserted into the viral backbone using the Gateway recombination system (Life Technologies). Briefly, antigen cassettes were cloned into an ENTRY plasmid, sequence-verified, and recombined *in vitro* with ChAdOx1-DEST or ChAdOx1-biDEST (containing a Gateway destination cassette in the E1 locus, or in both the E1 and the E4 loci, respectively). E1 and E4 expression cassettes both contained the TetR-repressible CMV promoter; E1 cassettes contained the BGH polyA sequence, while E4 expression cassettes contained the SV40 polyA sequence. E1 insertion occurs at the deleted E1 locus, while the E4 insertion site is located upstream of the intact E4 region.

Monovalent control vectors encode the vaccine antigen at the E1 locus.

Construction of the recombinant MVA vector was performed using BAC recombineering methods as described previously[13]. Briefly, EBOV and SUDV glycoprotein coding sequences (under the control of the short synthetic promoter (SSP) and the modified H5 (mH5) promoter, respectively) were inserted at the F11 locus, and MARV and LASV glycoprotein coding sequences (under the control of the short synthetic promoter (SSP) and the modified H5 (mH5) promoter, respectively) were inserted at the B8 locus.

### Virus production

Viral vectors were produced at the Viral Vector Core Facility at the Jenner Institute using standard methods[14] [12]. All adenovirus vectors were produced in the T-REx-293 cell line (Thermo Fisher Scientific), which allows for transcriptional repression of the vaccine antigens during vector production. Vectors underwent quality control (including titration, identity PCR and sterility testing) before being used in *in vitro* and *in vivo* studies.

### Expression testing

Expression of vaccine antigens from monovalent, bivalent and tetravalent viral vectors was assessed by western blot according to standard methods. Briefly, HEK293 cells were infected with vectors (MOI=1), harvested after 24 hours and lysed in RIPA buffer. Reduced and denatured lysates were resolved by 4-12% SDS-PAGE and transferred onto nitrocellulose. Glycoproteins were detected using mouse antiserum from mice previously vaccinated with monovalent vectors (ChAdOx1-EBOV, ChAdOx1-SUDV) or commercial anti-MARV GP antibody (ab190459, abcam) or anti-LASV GP2 antibody (ab190655, abcam), HRP-conjugated secondary antibody was added followed by chemiluminescence imaging (ChemiDoc, BioRad).

### Mouse immunogenicity studies

At least 6 week old female BALB/c mice (Envigo, UK) or CD-1 mice (Charles River, UK) were randomly distributed into individually ventilated cages on arrival, housed in groups of 3, 4, 5 or 6 under specific pathogen free conditions, fed and watered *ad libitum* with a 12:12 light-dark cycle.

In most cases, ChAdOx1-biEBOV and ChAdOx1-biLAMA were co-administered; mice received 50µl of ChAdOx1-biEBOV in the left rear leg and 50µl of ChAdOx1-biLAMA in the right rear leg. Monovalent mix refers to a mix of monovalent ChAdOx1-EBOV and ChAdOx1-SUDV administered in the left rear leg and a mix of monovalent ChAdOx1-MARV and ChAdOx1-LASV administered in the right rear leg. Blood samples (from the tail vein) were taken at various time points post vaccination. In prime-boost experiments, second vaccinations of 10^6^ PFU MVA expressing filovirus and LASV glycoproteins (‘tetraMVA’) were administered intramuscularly after the relevant time interval. Mice were culled humanely at the endpoint of the experiment via an approved Schedule 1 method; blood and spleens were harvested for further immunological analysis.

### ELISpot

Murine IFN-γ producing splenocytes were assessed by ELISpot assay after vaccination with filovirus viral vectors as previously described [15], with the following exceptions: Splenocytes were added to ELISpot plates at concentrations ranging from 1.25 x 10^5^ to 5 x 10^5^ cells/well and stimulated with pools of peptides at a final concentration of 1μg/mL per peptide. Peptide pools consisted of 15-mer peptides overlapping by 11 amino acids, spanning EBOV GP, SUDV GP, MARV GP or LASV GP. For graphical presentation, the number of IFNγ producing cells were calculated as the number of spot forming cells in the presence of peptides minus the number of spot forming cells without peptides and reported per million splenocytes.

### ELISA

Antibody responses were measured against trimerised EBOV GP (amino-acids 1-649 of GenBank protein AHX24649.1, with a C-tag), produced in house as described previously [11]. Antibody responses against monomeric SUDV GP (made in house) and recombinant MARV-Angola GP (Alpha Diagnostic International) were also measured. To test cross-reactivity against strains/species not in our vaccines, antibody responses were also measured against EBOV-Mayinga GP (Sino Biologicals), EBOV-Kikwit GP (Native Antigen Company), SUDV-Gulu GP (Sino Biologicals) and Bundibugyo (BDBV) GP (IBT Bioservices). Methods were as described previously[16] except that 2μg/ml of GP antigen was used to coat plates, blocking was performed with PBS/T containing 10% skimmed milk. Antibody responses against LASV were measured using pre-coated plates obtained from ReLASV® Pan-Lassa IgG/IgM ELISA Test Kit (GP) manufactured by Zalgen Labs, and all steps after blocking carried out as above. Reference pools of each of EBOV, SUDV, MARV and LASV antibody-positive mouse serum were used to form a standard curve for each plate. The relevant pool was added at an initial dilution of 1:250 (EBOV, MARV and LASV) or 1:125 (SUDV) in PBS/T and underwent ten 2-fold dilutions. An arbitrary number of ELISA units (AU) were assigned to the reference pool (62.5 AU for EBOV, MARV and LASV; 125AU for SUDV), and OD450 values of each dilution were fitted to a 4-parameter logistic curve using SOFTmax PRO software. ELISA units were calculated for each sample using the OD values of the sample and the parameters of the standard curve. All ELISA data presented are in AU, with the exception of IgG1/IgG2a ratios, which were calculated using OD.

### Intracellular cytokine staining (ICS)

Splenocytes were prepared as described above, plated in 96-well round bottom plates and stimulated using peptide pools for EBOV GP, SUDV GP, MARV GP or LASV GP (as described above) at a final concentration of 5µg/mL or media only. Stimulation and staining was then performed as described previously [17] except that the following antibodies were used: anti-CD4-Qdot605, anti-CD127-APCef780 (Invitrogen), anti-CD62L-PeCy7 anti-CD8-PerCP/Cy5.5 antibodies (eBioscience), LIVE/DEAD® Fixable Aqua Dead Cell Stain Kit (Thermo Fisher Scientific), anti-TNF-Alexa488, anti-IL-2-PE and anti-IFN-γ-e450 antibodies (eBioscience). Antigen-specific cells were identified by gating based on doublet negative, size, live cells and either CD4^+^ or CD8^+^ surface expression. Background responses in unstimulated control samples were subtracted from responses of peptide stimulated T cells.

### Measurement of cytokines and chemokines

Supernatants from ELISpot assays (as described above) were harvested after an 18-hour incubation and stored at −20⁰C. These samples were assayed using MSD Technology V-PLEX Mouse Cytokine 29-Plex kit according to the manufacturer’s instructions. Data analysis was performed using MSD Discovery Workbench 4.0. 18 of the 29 cytokines measured had detectable levels suitable for further analysis, which was performed using SPSS Statistics 25 (IBM) Correlations and heat maps were calculated and plots produced using RStudio (Version 0.99.903, packages corrplot (0.77) and package pheatmap).

### Guinea pig challenge experiment design

Groups of female Hartley strain guinea pigs (n=6/group) were intramuscularly vaccinated with 5 x 10^8^ IU of ChAdOx1-biEBOV or a mix of monovalent ChAdOx1 controls (ChAdOx1-EBOV and ChAdOx1-SUDV) or a negative control (ChAdOx1 with irrelevant antigen). 28 days after immunisation, the vaccinated animals were challenged subcutaneously with a lethal dose (10^3^ TCID_50_) of guinea pig-adapted EBOV (EBOV Yambuku-Ecran strain)[18]. The EBOV was passaged five times in guinea pigs to achieve lethality, as previously described[19]. Virus was titrated by 50% tissue culture infective dose (TCID50) assay in VeroE6 cells (European Collection of Cell Cultures, UK). Animals were assessed daily with respect to temperature and weight loss throughout the experiment. Clinical signs were monitored at least twice daily, and the following numerical score was assigned for analysis: 0 (normal); 2 (ruffled fur); 3 (lethargy, hunched and wasp waisted); 5 (rapid breathing); 10 (immobile, neurological). To prevent unnecessary suffering to animals, humane endpoints were used where animals would be culled upon reaching 10% weight loss and a moderate clinical sign or any of the following: 20% weight loss; immobility; paralysis; or neurological signs.

### IFNAR^-/-^ mouse challenge experiment design

Female IFNAR^-/-^ mice (A129, Marshalls BioResources, UK or #010830, Jackson Laboratory, USA) were immunised at the age of 6-8 weeks with 2 x 10^8^ infectious units (IU) of adenovirus vaccine in 50 µl via intramuscular injection in the right hind limb under anesthesia. Three weeks later mice were challenged with 3000 focus-forming units (ffu) or plaque-forming units (pfu) depending of virus in 100 ul via intraperitoneal injection under anesthesia. Four experiments were performed to assess efficacy of a single dose of biEBOV and biLAMA coadministered upon subsequent challenge with each of Ebola Zaire Guinea Kissidoguo, Ebola Sudan Boniface, Marburg Mukose and Lassa Josiah.

Within each experiment, controls receiving either an irrelevant ChAdOx1 (GFP) vector or the relevant monovalent control (ChAdOx1-EBOV, ChAdOx1-SUDV, ChAdOx1-MARV or ChAdOx1-LASV) was included. Six mice were vaccinated and challenged within each group, with the exception of ChAdOx1-GFP mice challenged with Marburg Mukose, for which there were only 5 mice. Mice challenged with EBOV, SUDV or MARV were assessed for up to 12 days after infection for signs of disease and weight loss and culled if reaching the humane end point. Based on previous studies of Lassa virus infection of IFNAR^-/-^ mice [20–22] 2 mice from each group (biEBOV+biLAMA, ChAdOx1-Lassa or ChAdOx1 GFP) were culled on day 4 and 8 with a further two to be culled on day 12 post-infection. However, on post-infection day 8, all mice in the control group had to be euthanized due to reaching the humane endpoint. The remaining mice in the two vaccine groups were spared until the end of the study (day 12).

### Statistics

Statistical analyses were carried out using GraphPad Prism version 7.01, unless otherwise stated. Grouped data are presented as means with SEM, unless otherwise indicated. Statistical significance of variations in continuous variables by group was analysed by Mann-Whitney or Kruskal-Wallis tests (for skewed data) or t-tests or ANOVA (for non-paired normally distributed data) as stated in results. For comparisons across multiple groups, Dunn’s multiple comparisons test was used for skewed data and Holm-Sidak multiple comparisons test was used for normally distributed data.

## Results

### Dual-antigen vaccine design

Two recombinant, dual-antigen ChAdOx1 vectors were constructed, one encoding SUDV GP at the E1 locus and EBOV GP at the E4 insertion site (ChAdOx1-biEBOV), and the other encoding LASV glycoprotein at the E1 locus and MARV glycoprotein at the E4 insertion site (ChAdOx1-biLAMA) (Fig 1a). Western blot analysis was performed to assess expression of the four antigens in infected cells. Cells infected with vectors expressing a single antigen only were used as positive controls (termed monovalent control vectors). Proteins of the correct size for all four inserts were expressed from ChAdOx1-biEBOV and -biLAMA (Fig 1b). We also generated a tetravalent MVA vector (tetraMVA) expressing all four glycoprotein antigens (Fig 1c) to test in a heterologous prime-boost regimen, as this was previously shown to elicit greater immunogenicity compared with homologous prime-boost or prime only, with either of the viral vector platforms [11] [17]. Western blot analysis of cells infected with the tetraMVA vector showed expression of all four antigens at levels equivalent to those of relevant monovalent controls (Fig 1d).

**Fig 1.**
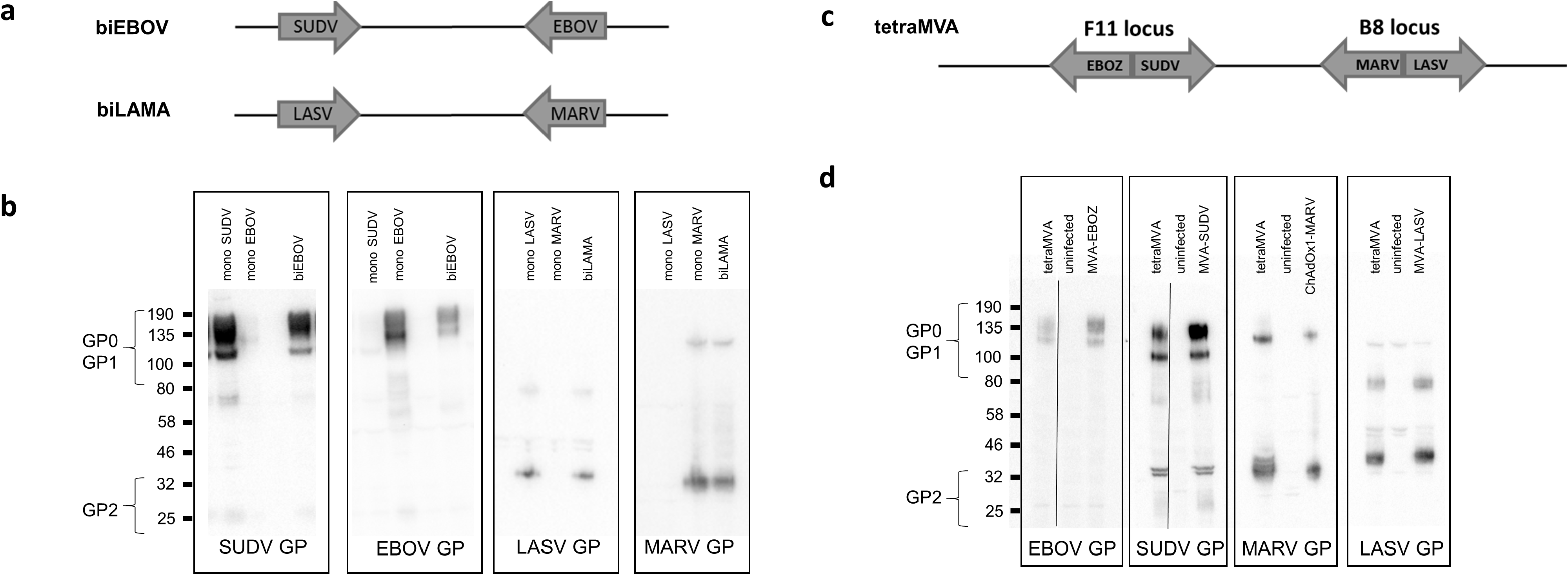
ChAdOx1 bivalent constructs. (a) Schematic to show design of bivalent vectors (b) Expression by western blot of each of the antigens in the bivalent constructs compared to monovalent controls (c) Schematic to show design of tetraMVA construct (d) Expression by western blot of each antigen in tetraMVA compared to monovalent controls. (S = SUDV GP, Z = EBOV GP, M = MARV GP, L =LASV GP)

### Optimisation of administration regimen

We and others have previously observed differences in cellular immunogenicity when two monovalent vaccines are administered compared to a single bivalent vaccine that encodes the same two antigens [23]. We investigated the difference in immunogenicity between administration of two separate vaccine constructs encoding monovalent antigens with a single construct that encodes both (biEBOV versus monovalent mix and the biLAMA versus monovalent mix). Very minimal differences in cellular immunogenicity to either antigen was observed whether two monovalent vaccines, or a single bivalent vaccine, was administered (Suppl Table 1).

However, when combining both bivalent vaccines (biEBOV and biLAMA) into a single shot and comparing it to a mix of all four monovalent vaccines in a single shot, lower T cell responses were measured against the EBOV antigen (Suppl Table 1 – regimen 3). We have previously found that the administration of the two competing vectors into different muscles alleviated the cellular immune competition and can improve immunogenicity [23]. Therefore, we administered biEBOV and biLAMA to the left and right rear leg, respectively and compared this to a mix of monovalent EBOV and SUDV vaccines (left leg) and a mix of monovalent LASV and MARV vaccines (right leg). This improved the response of the bivalent, in comparison to monovalent, regimens (Suppl Table 1 - regimen 4). Co-administration was, therefore, considered the optimal regimen going forward and was used unless otherwise stated.

### Humoral immunogenicity

Next, we assessed the humoral immune response after co-administration of our bivalent vaccines; individual dual-antigen vaccines were delivered to a separate hind leg of a mouse (e.g. biEBOV to the right hind limb and biLAMA to the left hind limb). For comparison, the monovalent controls were mixed as follows; ChAdOx1-EBOV and -SUDV were mixed and administered to one leg, and ChAdOx1-MARV and -LASV were mixed and administered to the other leg. 6 weeks post vaccination, humoral immune responses were assessed by ELISA against each of the four antigens in the outbred CD-1 mouse strain. Across all four antigens, there were no significant differences (Mann-Whitney test) in the total IgG titres elicited by the dual-antigen vaccines compared to the mixed monovalent controls (Fig 2a).

**Fig 2.**
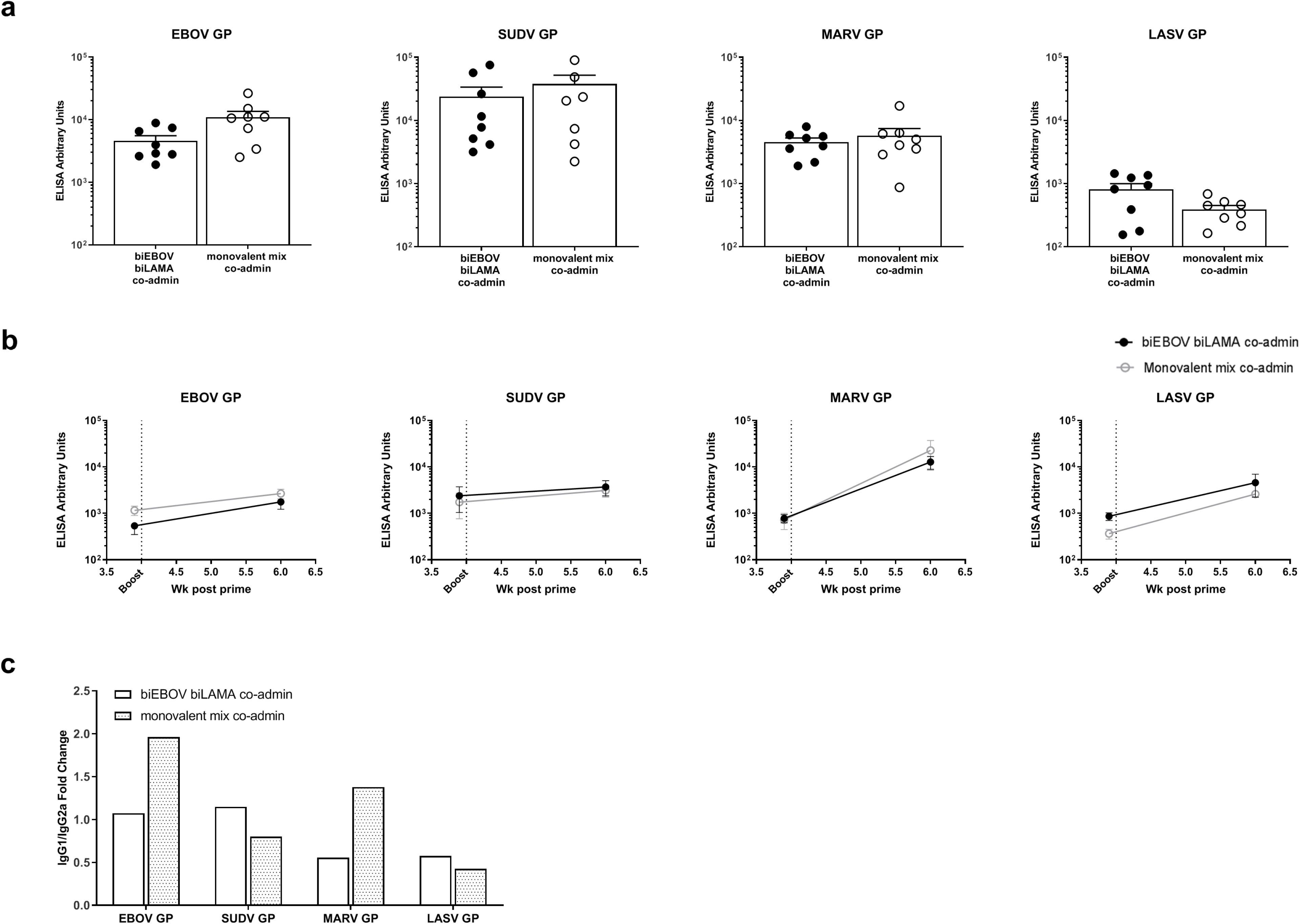
Antibody levels induced by ChAdOx1 dual antigen prime, and following tetraMVA boost. (a) CD-1 mice were immunised with either biEBOV and biLAMA co-administered or a mix of monovalent controls. Total IgG levels were measured 6 weeks later. (b) BALB/C mice were primed with either biEBOV and biLAMA co-administered or a mix of monovalent controls, 4 weeks later all mice were boosted with tetraMVA. Total IgG levels were measured pre-boost and at 2 weeks post boost vaccination. (c) IgG1 and Ig2a isotype levels were measured in CD-1 mice 8 weeks after prime vaccination, and are expressed as a ratio.

To augment humoral immunity, BALB/c mice that had been primed with the dual-antigen ChAdOx1 vaccines were boosted with the tetraMVA vector expressing all four antigens. The humoral immune response to the four antigens was increased at two weeks post-boost (6 weeks post-prime) compared to pre-boost levels to varying degrees (Fig 2b).

To further delineate antibody responses, we performed isotype-specific ELISAs. Initially, this was performed in the inbred BALB/c strain and the relative Th1/Th2 contributions were evaluated as the ratio of IgG1/IgG2a (Suppl Fig 2). After prime vaccination with bivalent vaccines, the antibody responses were approximately Th1-Th2 balanced for all 4 antigens. After boost vaccination with tetraMVA, the response against MARV GP was Th2-biased while responses remained balanced against the other three antigens. To account for potential bias introduced by a fixed haplotype in the BALB/c model, we also measured IgG1 and IgG2a in CD-1 mice. We detected similar levels for IgG1 and IgG2a in CD-1 mice vaccinated with our bivalent vectors and a mix of monovalent controls (no significant differences, Mann-Whitney test). In CD-1 mice vaccinated with bivalent vaccines, the IgG1/IgG2a ratios against MARV GP and LASV GP antigens suggest skewing towards Th2, whereas the ratios for EBOV GP and SUDV GP were Th1-Th2 balanced (Fig 2c). These combined results demonstrate that the humoral immune response (Th1 or Th2) after vaccination with our multi-antigen vectors differ between mouse strains, can vary after prime and prime-boost vaccination and may be antigen-rather than vector-dependent. The levels of other isotypes (IgG2b, IgG3, IgM and IgA) were assessed, but were below the limit of detection in our assay.

It will be important for new vaccines against VHF to be protective across variant viral strains. To measure humoral reactivity to different filovirus isolates, CD-1 mice were primed with co-administered bivalent ChAdOx1 vectors or a mix of monovalent controls. Cross-strain humoral immunity was measured against the following filovirus glycoproteins: EBOV-Mayinga GP (AAC54887.1), EBOV-Kikwit GP (048.1), SUDV-Gulu GP (YP_138523.1) and BDBV GP (ACI28624.1). Total IgG levels 6 weeks after vaccination demonstrated that antibody responses against all variant proteins could be detected (Suppl Fig 2a). Booster vaccination with MVA has previously been shown to not only increase but also broaden antigen-specific immune responses [24]. To assess if the cross-reactive immune responses could be augmented, bivalent-primed BALB/c mice were boosted with tetraMVA 4 weeks post-prime. At 6 weeks post-boost, total IgG antibody titres increased (by at least 2-fold) against all variant filovirus strains except for SUDV Gulu (Suppl Fig 2b). In all cases, responses to the bivalent vaccines were comparable to the monovalent control mix. Naïve mouse serum did not react with the antigens in the assays.

### Cellular immunogenicity

Immunogenicity can vary in different mouse strains so both CD-1 and BALB/c mice were immunised with biEBOV and biLAMA vaccines to assess the cellular immunogenicity of all four VHF antigens. The bivalent vaccines were co-administered, and immunogenicity was assessed by ELISpot 2 weeks post vaccination (Fig 3a and b). In CD-1 mice (Fig. 3a), the magnitudes of ELISpot responses were comparable between the co-administered dual-antigen ChAdOx1 vectors and the monovalent controls for EBOV GP and MARV GP. However, for SUDV GP and LASV GP, the ELISpot responses were significantly higher in the dual-antigen regimen (p = 0.013 and p = 0.007 respectively, Mann-Whitney test). In BALB/c mice (Fig. 3b), there were no differences in the ELISpot responses between the two regimens for SUDV GP, MARV GP and LASV GP, but the response to EBOV GP was significantly higher following vaccination with co-administered dual-antigen vaccines than for the monovalent controls (p = 0.0014, Mann-Whitney test).

**Fig 3.**
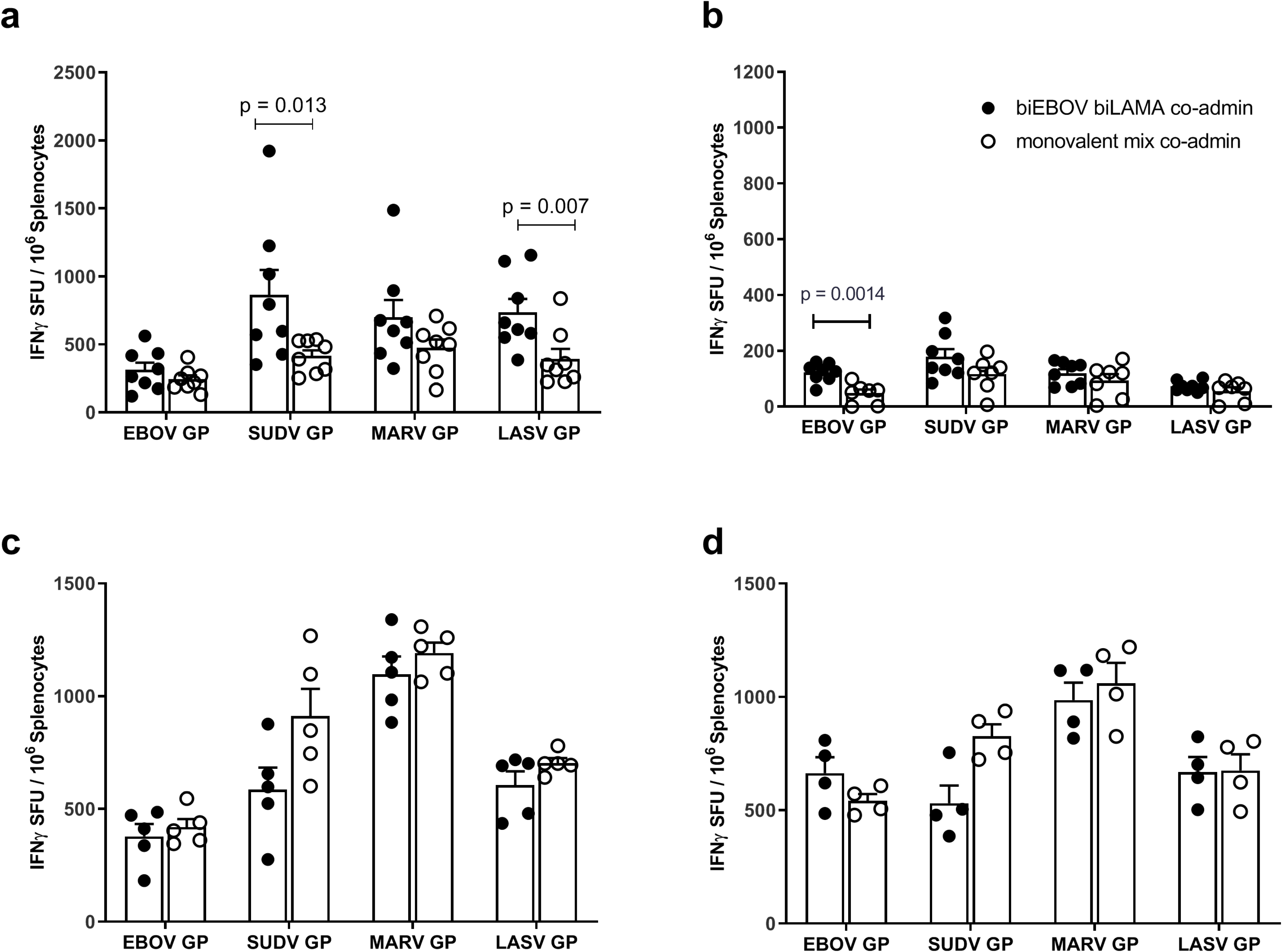
T cell immunogenicity of ChAdOx1 dual antigen prime and prime-boost (ChAdOx1 and MVA) approaches. In all panels mice received either biEBOV and biLAMA co-administered or a mix of monovalent controls. IFNγ ELISpot in CD-1 mice 8 weeks post vaccination (a) and in BALB/C mice 2 weeks post vaccination (b). In experiments shown in (c) and (d), BALB/C mice were primed as described above and 4 weeks later boosted with tetraMVA. IFNγ ELISpot 2 weeks post boost vaccination (c) and 8 weeks post boost vaccination (d), *p* values determined by Mann-Whitney tests. X-axis labels show peptide pools used for splenocyte stimulation.

Cellular immunogenicity of a prime-boost regimen was assessed in BALB/c mice at 2 weeks and 8 weeks after a tetraMVA boost (Fig 3c and 3d). No statistically significant differences were observed between the co-administered dual-antigen ChAdOx1 vectors and monovalent mix control for the four antigens, although a trend toward lower responses in the dual-antigen group was observed for SUDV GP. ELISpot responses in both groups were substantially boosted 2 weeks after tetraMVA administration (Fig 3b and 3c). Notably, SUDV, MARV, and LASV GP ELISpot responses did not wane from week 2 to 8, and EBOV GP-specific responses further increased during this period (Fig 3c and 3d).

Intracellular cytokine staining was performed on splenocyte samples harvested 2 weeks after prime vaccination (Fig 4a), and 2 weeks after tetraMVA boost (Fig 4b). Antigen-specific responses primarily consisted of IFNγ^+^ CD8^+^ T cells, the majority of which also secreted TNFα; a small proportion of these cells additionally produced IL-2 (Fig 4, bottom panels). The highest percentage of cytokine positive CD8^+^ T cells were detected against SUDV GP, followed by LASV GP. Meanwhile, the lowest responses were measured against EBOV GP. For CD4^+^ T cells, MARV GP-specific responses demonstrated the highest percentage of cytokine positive CD4^+^ T cells and the lowest responses were measured against EBOV GP (Fig. 5). Of the vaccine-specific CD4^+^ responses post prime, there was a greater percentage of triple cytokine positive cells detected compared to cells that were double positive for IFNγ and TNFα against EBOV GP, SUDV GP, MARV GP and LASV GP (Fig. 5c). These data imply that the hierarchy of immune dominance of VHF antigens responses may differ depending on the response (CD8^+^ and CD4^+^) being measured.

**Fig 4.**
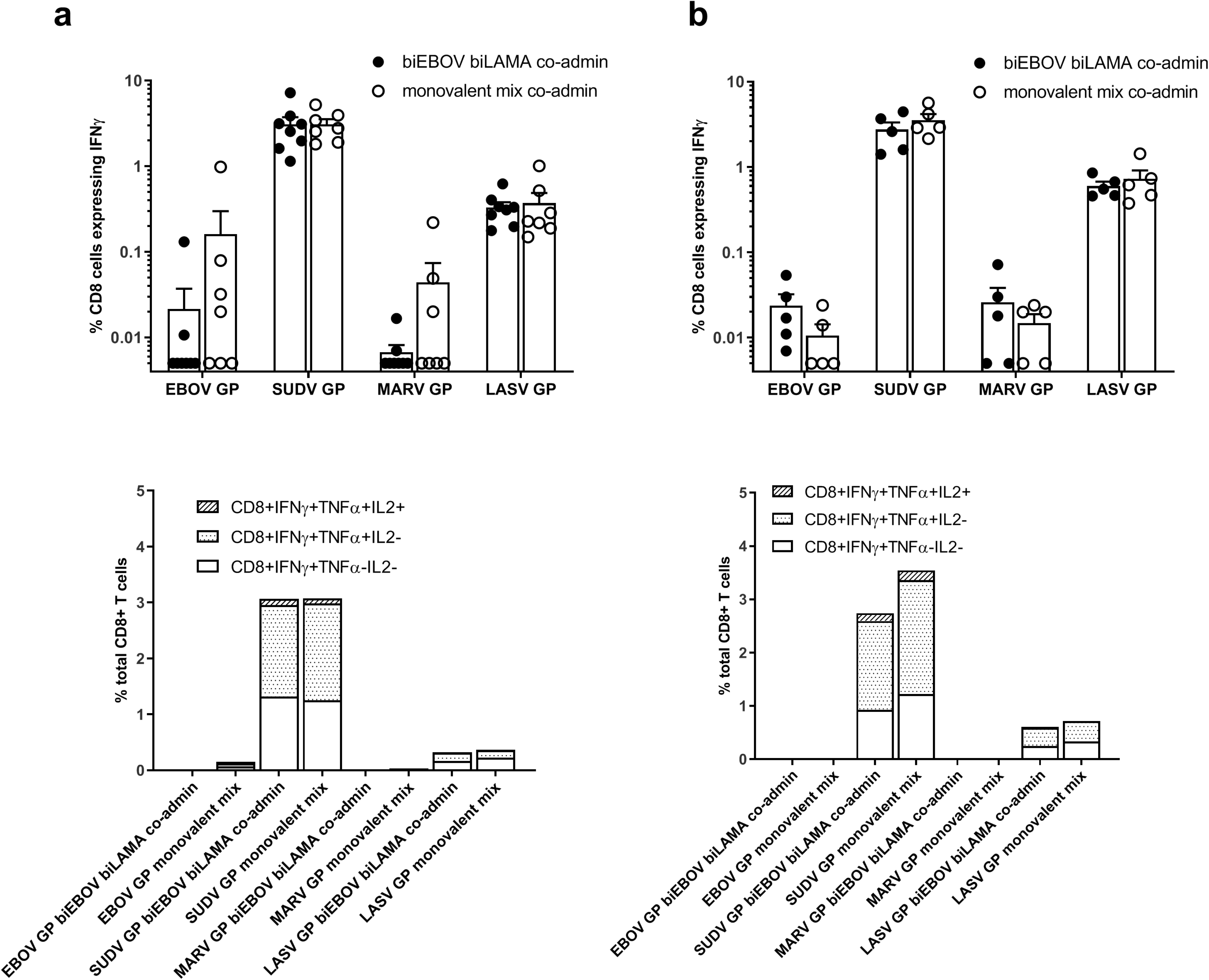
CD8+ T cell immunogenicity of prime only and prime-boost vaccination determined by ICS. BALB/c mice were primed with either biEBOV and biLAMA co-administered or a mix of monovalent controls, and in one experiment (b) mice were boosted with tetraMVA 4 weeks later. Cytokine levels determined by ICS (a) 2 weeks post prime, (b) 2 weeks post boost Top panels – IFNγ^+^ CD8^+^ T cells. X-axis labels in top panels show peptide pools used for splenocyte stimulation. Bottom panels – CD8^+^ T cells expressing IFNγ, TNFα and IL-2

**Fig 5.**
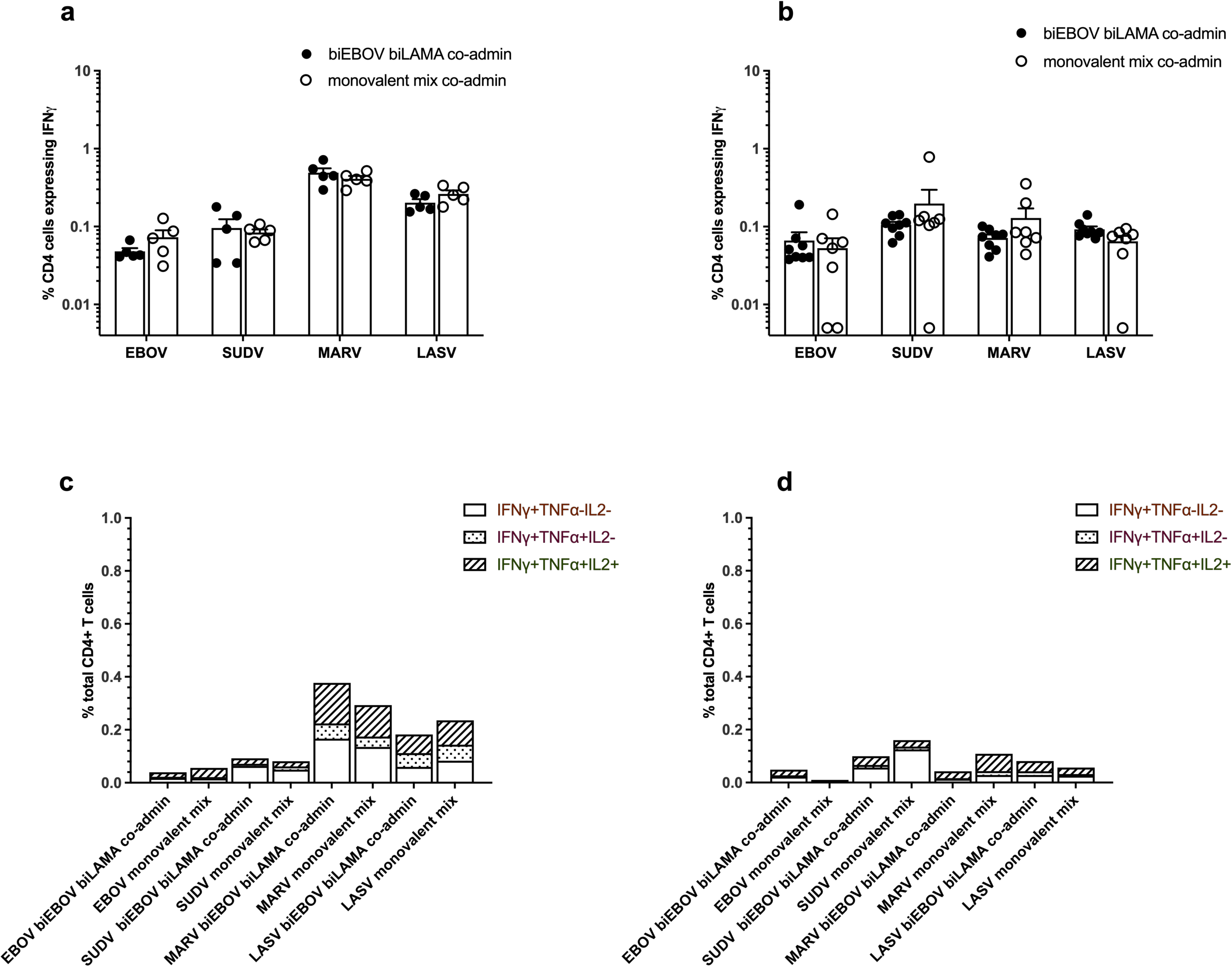
CD4+ T cell immunogenicity of prime only and prime-boost vaccination determined by ICS. BALB/c mice were primed with either biEBOV and biLAMA co-administered or a mix of monovalent controls, and in one experiment (b) mice were boosted with tetraMVA 4 weeks later. Cytokine levels determined by ICS (a) 2 weeks post prime, (b) 2 weeks post boost Top panels – IFNγ^+^ CD4^+^ T cells. X-axis labels in top panels show peptide pools used for splenocyte stimulation. Bottom panels – CD4^+^ T cells expressing IFNγ, TNFα and IL-2

A multiplex mouse cytokine kit was used to measure antigen-specific vaccine-induced expression of 28 different cytokine/chemokines from splenocyte samples 2 weeks after boost vaccination (sample sets were as analysed in Fig 4b). Advantageously, the multiplex assay can measure cytokines not routinely measured by ELISpot or ICS and can generate a more complete picture of the immune profile post-vaccination. Only those cytokines and chemokines that could be reliably measured post vaccination were tested for clustering - 18 different cytokines were included (Suppl Fig 3). The data indicate that groups of certain cytokines are more likely to be concurrently detected e.g. IL-1β, TNFα, IP-10, IFNγ, IL-2, IL-5, IL-4, IL-10 and IL-6 were clustered together and the induction of these cytokines following stimulation with filovirus peptides is to be expected[25]. Cytokine fold change (when compared to unstimulated cells) was calculated for each of the four antigens tested (Suppl Table 1) and interactions between cytokines across samples were depicted in a heatmap (Fig 6). Across all four antigens, IFNγ was the most highly expressed cytokine, followed by MIP-1α, IL-17 and MIP-2. IL-6, IL-4, IL-2, IL-10 and IL-5 were also highly expressed in supernatant from MARV GP-stimulated cells. Supernatant from SUDV GP-stimulated cells displayed a different pattern of cytokine expression. A number of cytokines were produced at lower levels by SUDV GP stimulated cells compared to unstimulated. However, no significant differences in fold change between the monovalent mix and dual-antigen vaccination regimens were observed when stimulated with EBOV, SUDV or MARV GPs (Suppl Table 1).

**Fig 6.**
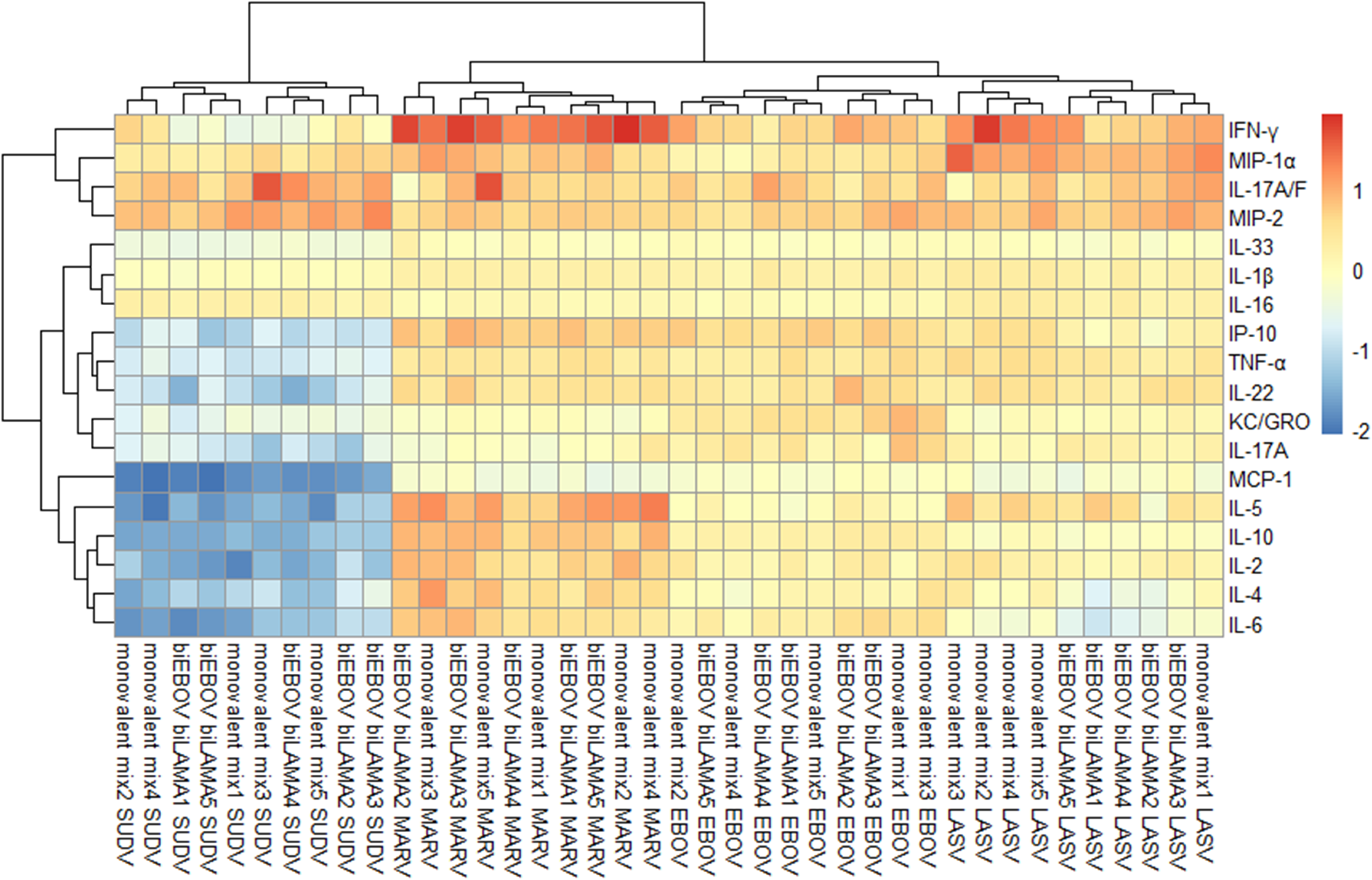
Cytokine and chemokine analysis after prime-boost vaccination. BALB/c mice were primed with either biEBOV and biLAMA co-administered (n=5) or a mix of monovalent controls (n=5), 4 weeks later all mice were boosted with tetraMVA. Cytokine levels (in supernatant from splenocytes stimulated with the peptide pools for EBOV GP, SUDV GP, MARV GP and LASV GP) determined by MSD technology platform 2 weeks post boost. Heat map shows fold change in peptide stimulated vs unstimulated supernatant across peptide pools (each stimulation for splenocytes from each mouse is labelled 1-5) and vaccine regimen. Red indicates positive fold change, blue indicates negative fold change.

Overall, humoral and cell-mediated (ELISpot, flow cytometry and cytokine) immunogenicity data demonstrated that the dual-antigen vaccines can induce immune responses, and that these responses are diverse, long-lived and polyfunctional.

### Lethal challenge to assess protection

We next assessed the efficacy of our biEBOV vaccine in a lethal guinea pig EBOV challenge model (using a guinea pig-adapted EBOV based on the EBOV-Yambuku-Ecran strain, GenBank: AF086833.2) (Suppl. Fig 4a). Guinea pigs vaccinated with ChAdOx1-biEBOV or a mix of ChAdOx1-EBOV and -SUDV controls survived the duration of the challenge, whereas all control animals succumbed to infection and reached the humane end point by day 7-9 post-infection (Suppl. Fig 4b). Animals vaccinated with the control ChAdOx1 GFP vaccine lost weight from day 5 onwards, and exhibited an increase in temperature (Suppl. Fig 4c,d). Animals vaccinated with biEBOV or a mix of monovalent control vaccines continued to gain weight post challenge with no significant temperature fluctuations (Suppl. Fig 4c,d). During the study, guinea pigs were clinically assessed; clinical signs were first observed in the ChAdOx1 GFP negative-control group on day 5 post challenge (Suppl. Fig 4e). No animal vaccinated with biEBOV or a mix of monovalent controls exhibited any clinical signs (Suppl. Fig 4e).

To further assess efficacy of biEBOV and biLAMA we utilised IFNAR^-/-^ mice, due to their increased susceptibility to infection, for EBOV, SUDV, MARV and LASV virus challenges. For each virus, mice were vaccinated with either a co-administration of biEBOV and biLAMA, the monovalent controls or the irrelevant control and subsequently infected with specific virus as described (Fig 7a). In the control groups (ChAdOx1 GFP), 50% and 100% of the mice infected with EBOV or SUDV, respectively, succumbed to infection (Fig 7b,c). In contrast, a single dose of biEBOV and biLAMA co-administered resulted in protection from EBOV (Fig 7b) and SUDV (Fig 7c) infection (p = 0.028 and p<0.001 respectively). Similar results were observed in mice infected with Marburg Mukose although this did not reach significance as the challenge was not uniformly lethal (Fig 7d) (p = 0.068). Disease was reflected in weight loss and clinical signs such as piloerection, decrease in activity and general condition, while animals receiving vaccination with either biEBOV and biLAMA or monovalent control lost none or very little weight and showed no clinical signs (Suppl Fig 5a to d). Complete protection against Lassa virus infection due to biEBOV + biLAMA vaccination was not observed in this study (Fig 7e and supplemental fig 5d).

**Fig 7.**
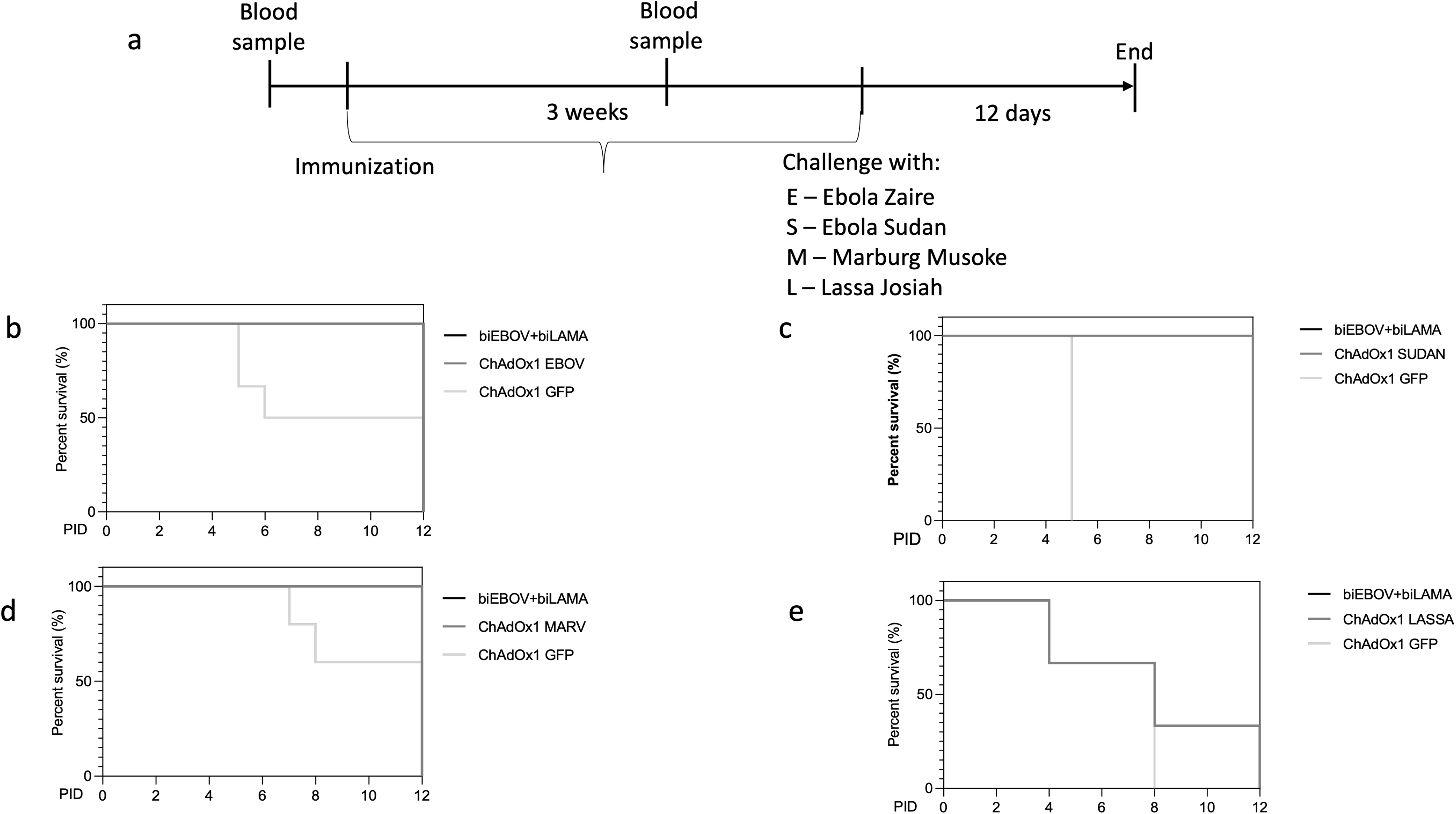
Efficacy of biEBOV and biLAMA against EBOV, SUDV, MARV and LASV infection in mice. Groups of 6 IFNAR^-/-^ mice received either biEBOV and biLAMA, relevant monovalent control or irrelevant control immunizations and were challenged 3 weeks later via i.p injection of specific virus. Animals were culled upon reaching the humane end point or the end of the study (12 days post-infection) (a) Experimental Design. Survival analysis for each of the challenge viruses: (b) Ebola Zaire Guinea Kissidoguo, (c) Ebola Sudan Boniface (d) Marburg Mukose and (e) Lassa Josiah. p values determined by Log rank (Mantel-Cox) test

## Discussion

This work addresses the global need for vaccines in preparedness for future outbreaks of haemorrhagic fevers caused by EBOV, SUDV, MARV and LASV. The unprecedented ease of EVD spread in West and Central Africa in the preceding 20 years along with the recent MARV and SUDV outbreaks in Tanzania, Equatorial Guinea, and Uganda, exemplify the consequences of having no deployable prophylactic intervention against a high-impact outbreak pathogen. Indeed, the geographical overlap of *Filoviridae* and LASV continues to expand. This expanding overlap is partially due to better detection methods, including the isolation of genetically diverse filoviruses from flying fruit bats (e.g. *Rousettus aegyptiacus*) and the identification of new hosts for LASV [26–28]. The large geographical areas habituated by these natural hosts, which can travel vast distances, indicate that there is an increased risk of outbreaks in previously unaffected areas and likely an increased incidence of undocumented subclinical infection [26–30].

During the 2013-2016 West African EBOV outbreak, six of seven vaccine candidates tested in clinical trials employed viral vector technologies, including three adenoviral and two MVA vectored vaccines. Results from the Phase III rVSV-ZEBOV clinical trial run during the 2013-2016 EBOV outbreak demonstrated that a single-dose regimen of a viral vectored vaccine (rVSV-ZEBOV) is both safe and efficacious in preventing EVD [8]. While correlates of protection are not clearly defined for all VHF, a protective role for antibodies has been demonstrated in EBOV [31–33]. Comparison of the protective neutralising antibody responses achieved after rVSV-ZEBOV immunisation with those elicited by an adenoviral vectored EBOV vaccine candidate showed that the latter induced equivalent antibody levels [11], providing encouraging evidence to support further clinical development of adenoviral vaccine vectors against viral haemorrhagic fevers. In mice, neutralising antibodies are typically IgG1, while IgG2a antibodies are associated with induction of stronger ADCC and complement activation [34]. Overall, our data indicate that the bivalent vaccines induce a mixed humoral response inducing both IgG1 and IgG2a post-vaccination. The levels of other isotypes (IgG2b, IgG3, IgM and IgA) were assessed but were below the limit of detection in our assay. We also demonstrate that immunisation with bivalent vaccines induces antibodies which recognise proteins from different isolates of filoviruses. Further work will be performed to demonstrate the breadth of this heterosubtypic immunity.

Importantly, the adenoviral vectored vaccines tested here also induce high levels of T cell immunity which have been demonstrated to play an important role during VHF outbreaks caused by EBOV and LASV in particular (reviewed [35]). Ebolavirus challenge studies in recombinant adenovirus– vaccinated macaques have demonstrated that protection is associated with robust dual-cytokine secreting (IFNγ and TNFα) CD8^+^ T cells[36] [37]. Depleting these macaques of CD8^+^ T cells before challenge abolished vaccine-mediated protection [38], suggesting that CD8^+^ T cells, and/or CD8^+^ NK cells, are necessary for virus clearance in this model system. Encouragingly, the predominant cellular immune response observed following vaccination with our bivalent vaccines was IFNγ^+^ CD8^+^ T, with a large proportion also being TNFα^+^. Our multiplex cytokine data showed an array of cytokines induced by antigenic peptides after filovirus and arenavirus vaccination and adds breadth to the customarily reported IFNγ, TNFα and IL-2 expression in T cells. We observed some differences in the cytokine profile of cells stimulated with SUDV peptides compared to the other three antigens. Of note, IFNγ was not as highly expressed as for the other three antigens. In contrast, ELISpot and flow cytometry data showed that SUDV peptide stimulation elicited higher numbers of IFNγ producing cells. However, the differential stimulation times with SUDV peptides may exhaust the cells or be cytotoxic, resulting in lower levels of IFNγ being detected in supernatant and measured in our multiplex cytokine assay.

Importantly, we have demonstrated that the two bivalent vaccines perform well in both challenge and immunogenicity experiments. We also demonstrate here that a single dose of our biEBOV and biLAMA vaccines co-administered can confer protective efficacy in a lethal challenge model for three of our viruses of interest (EBOV, SUDV, and MARV), but not LASV. Further work is needed to explore these findings, as none of the animals who were vaccinated with a LASV GP regimen reached their humane endpoint while the controls all reached theirs by day 8. In addition, we have previously demonstrated that NHPs (unpublished data) and guinea pigs immunised with a single dose of ChAdOx1-LASV alone was protective against challenge [39]. This difference in challenge outcome may be due to differences in animal challenge models or potential interference associated with administration of multiple vaccines concurrently.

While short-lived but efficacious immunity following a one-shot vaccination regimen may be sufficient for curbing a pandemic, long-lasting immunity is considered a pre-requisite for first-line responders. A heterologous prime-boost vaccine regimen could be particularly appropriate for healthcare practitioners and first-line responders, as this regimen is demonstrated here and in clinical trials to induce long-lived, high-titre responses that will be best-placed to protect against repeated high-risk exposure to the viral targets. Chimpanzee Adenovirus (ChAd)-based vaccines are frequently administered as the first highly immunogenic vaccination (prime) and can be followed with a second, MVA-based vaccination (boost) – this ChAd-MVA ‘prime-boost’ regimen benefits from remarkable and long-lived immunogenicity [4]. For example, the longevity of high titre neutralising antibodies and cellular immune responses to EBOV GP has been demonstrated following a ChAd-MVA prime-boost regimen both in man and macaques, where protective efficacy, in the latter, against EBOV challenge was significantly extended post-vaccination following a boost immunisation [36] [11] [40]. In addition to our dual-antigen adenoviral vectors, we therefore also developed a multi-antigen MVA vector that encodes all four VHF antigens (tetraMVA). Using tetraMVA in a prime-boost regimen together with the adenoviral vectors induced a higher cellular and humoral response and a diverse chemokine/cytokine profile.

In summary, using both single-shot (for rapid protection) and prime-boost (for long-lived protection) regimens will allow for optimal outbreak pathogen control, to curb epidemics and avoid future public health and humanitarian crises. Further clinical development of these vectors will contribute to the common goal: deployable vaccine solutions for future VHF outbreaks.

## Acknowledgements

We would like to thank the excellent animal handling staff of Biomedical Sciences (BMS), University of Oxford and the Biological Investigations Group (BIG), UKHSA (formerly Public Health England) along with the UKHSA (PHE) High Containment Microbiology group (HCM) for assistance of work at Containment Level 4. This work was funded by Innovate UK (Novel multivalent vaccines against haemorrhagic fevers, 971510) and MRC (Confidence in Concept CiC 2015-16, MC_PC_15040, Liverpool School of Tropical Medicine). Views expressed are those of the authors and do not necessarily reflect those of the employing institutions. In addition, we would like to thank the staff at Astrid Fagræus laboratorium, Karolinska Institute for their excellent help with sampling and animal care. In addition, we would like to thank the staff at Astrid Fagræus laboratorium, Karolinska Institute for their excellent help with sampling and animal care.

---

**Suppl Fig 1:**
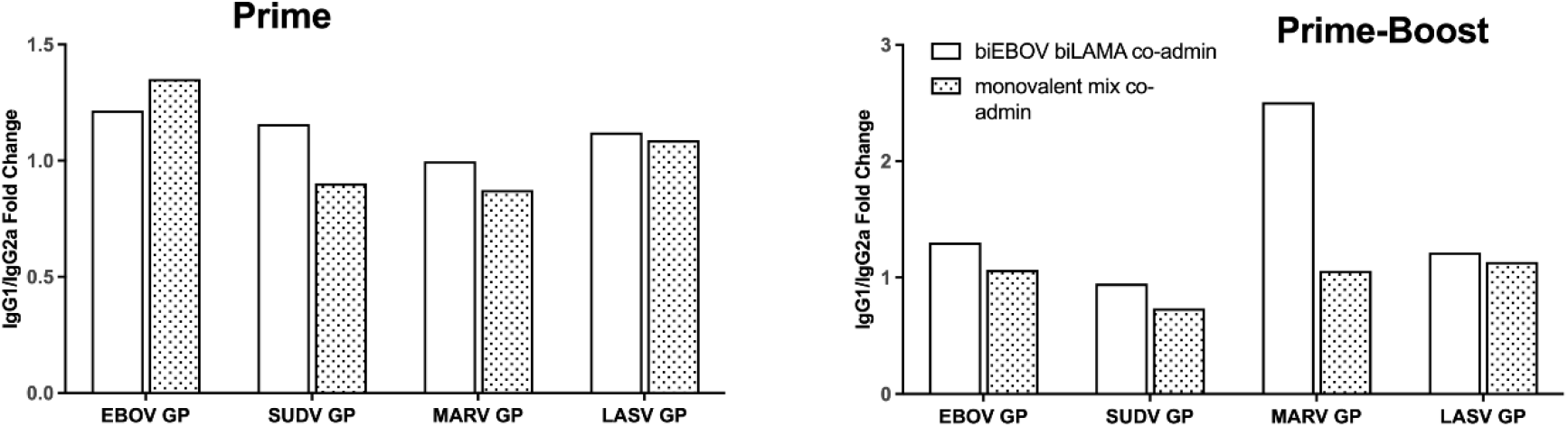
Antibody isotype levels induced by ChAdOx1 dual-antigen prime and tetraMVA boost. BALB/c mice were primed with either co-administered biEBOV and biLAMA or a mix of monovalent controls and boosted with tetraMVA 4 weeks later. IgG1 and Ig2a isotype levels are expressed as a ratio at 4 weeks after prime vaccination (left panel) and 8 weeks after boost vaccination.

**Suppl Fig 2:**
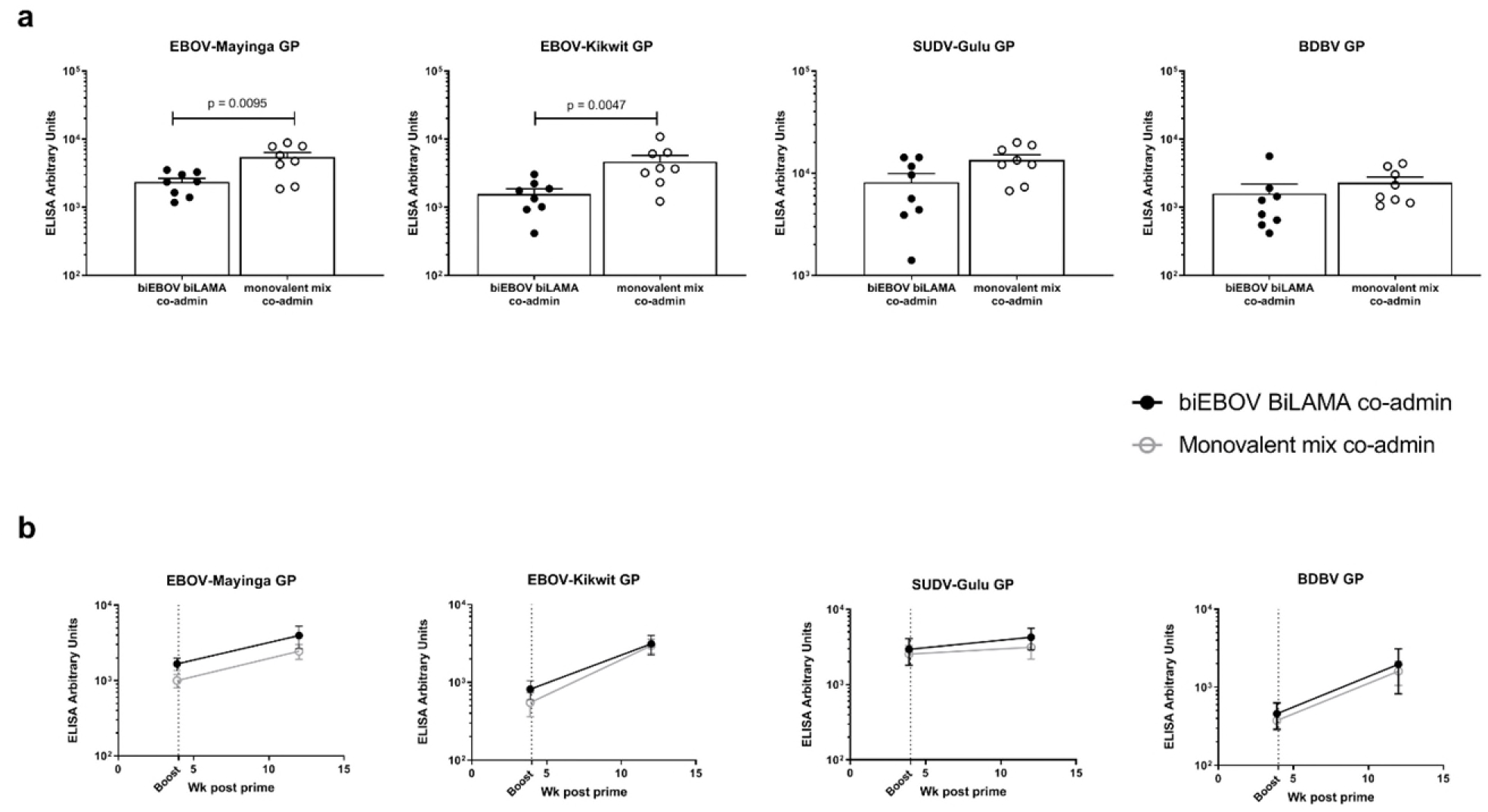
Cross reactivity induced by ChAdOx1 dual antigen prime and followed by tetraMVA boost. Antibody titres against EBOV-Mayinga GP, EBOV-Kikwit GP, SUDV-Gulu GP and BDBV GP were assessed. (a) CD-1 mice were primed with either biEBOV and biLAMA co-administered or a mix of monovalent controls. Total IgG levels were measured 8 weeks later (p values determined by Mann Whitney test). (b) BALB/c mice were primed with either biEBOV and biLAMA co-administered or a mix of monovalent controls. 4 weeks later all mice were boosted with tetraMVA. Total IgG levels before boost and at 8 weeks post boost vaccination.

**Suppl Fig 3:**
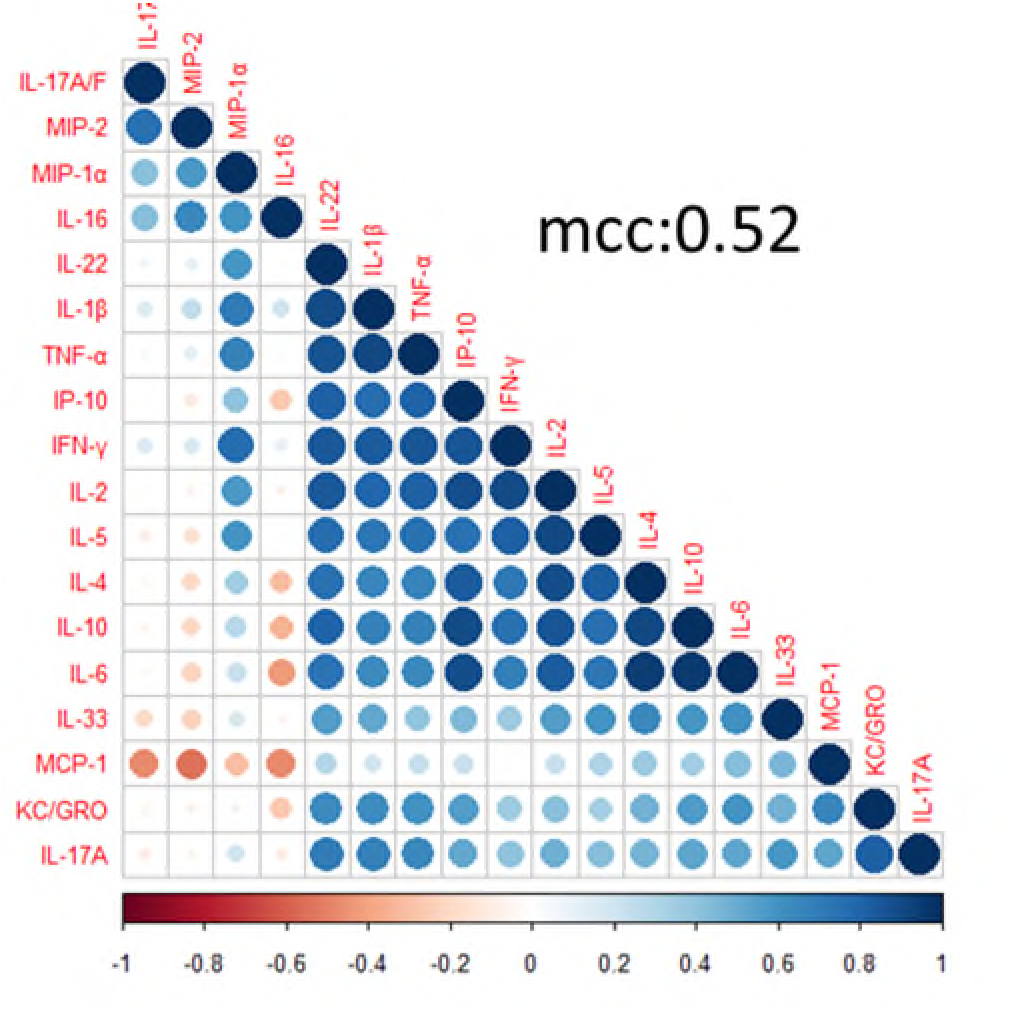
Interaction of cytokines and chemokines after prime-boost vaccination. Mice were primed with either biEBOV and biLAMA co-administered or a mix of monovalent controls. 4 weeks later, all mice were boosted with tetraMVA. Cytokine levels (in the supernatant of peptide pool-stimulated splenocytes) at 2 weeks post boost were measured using a commercial assay (MSD). Correlation matrix among 18 cytokines measured, red indicates negative correlation, blue indicates positive correlation.

**Suppl Fig 4:**
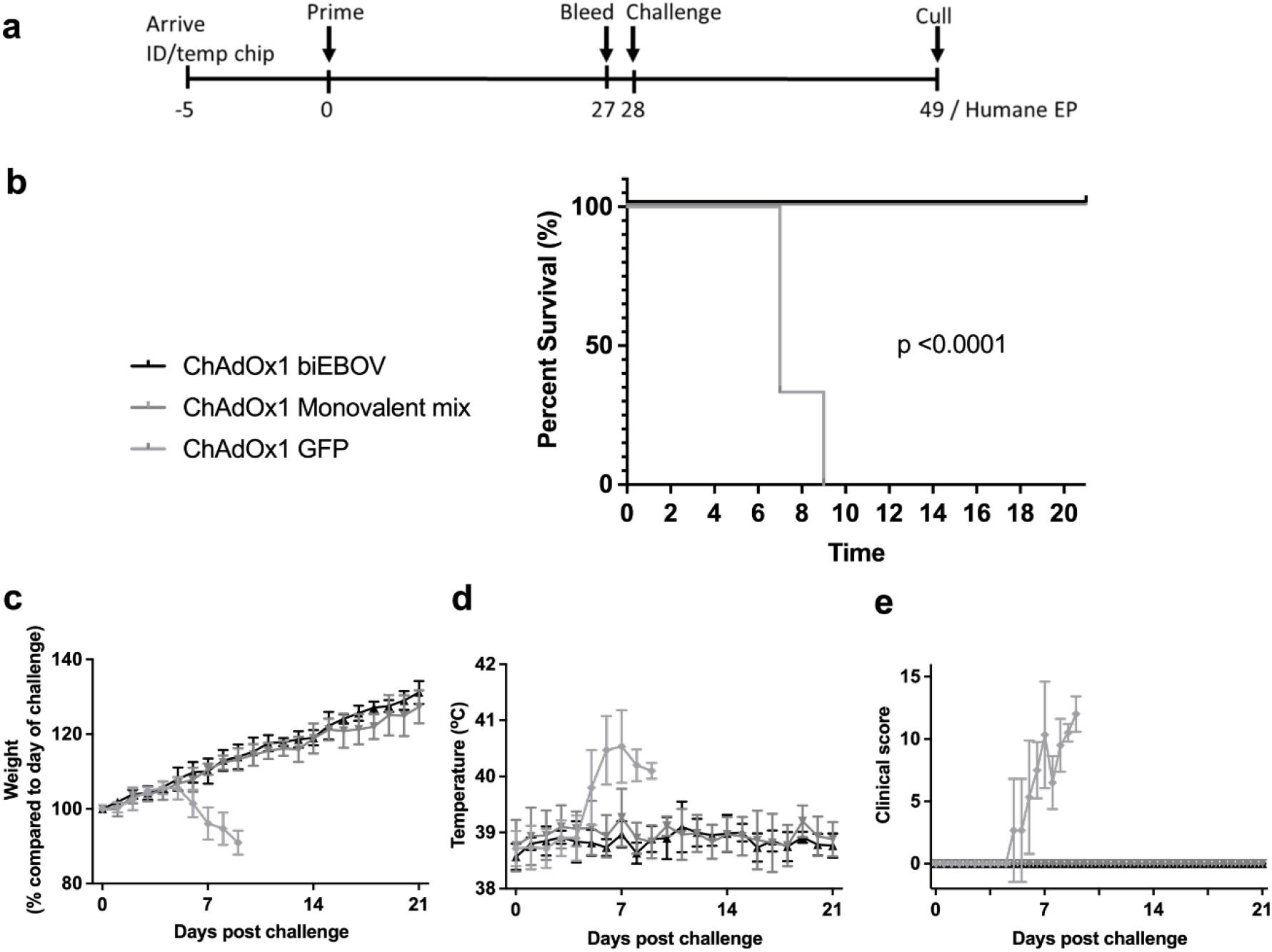
Protective efficacy of ChAdOx1 bivalent vaccines against Ebola Zaire infection in Guinea Pigs. Groups of 6 Hartley Guinea Pigs received either biEBOV, mix of monovalent controls or control without antigen and were challenged 4 weeks later IP with Ebola virus. Animals were culled upon reaching humane end point or 21 days post challenge. (a) Experimental design (b) Survival analysis. *p* value determined by Log rank (Mantel-Cox) test (c) Weight change post challenge (d) Temperature post challenge (e) Clinical score post challenge

**Suppl Fig 5.**
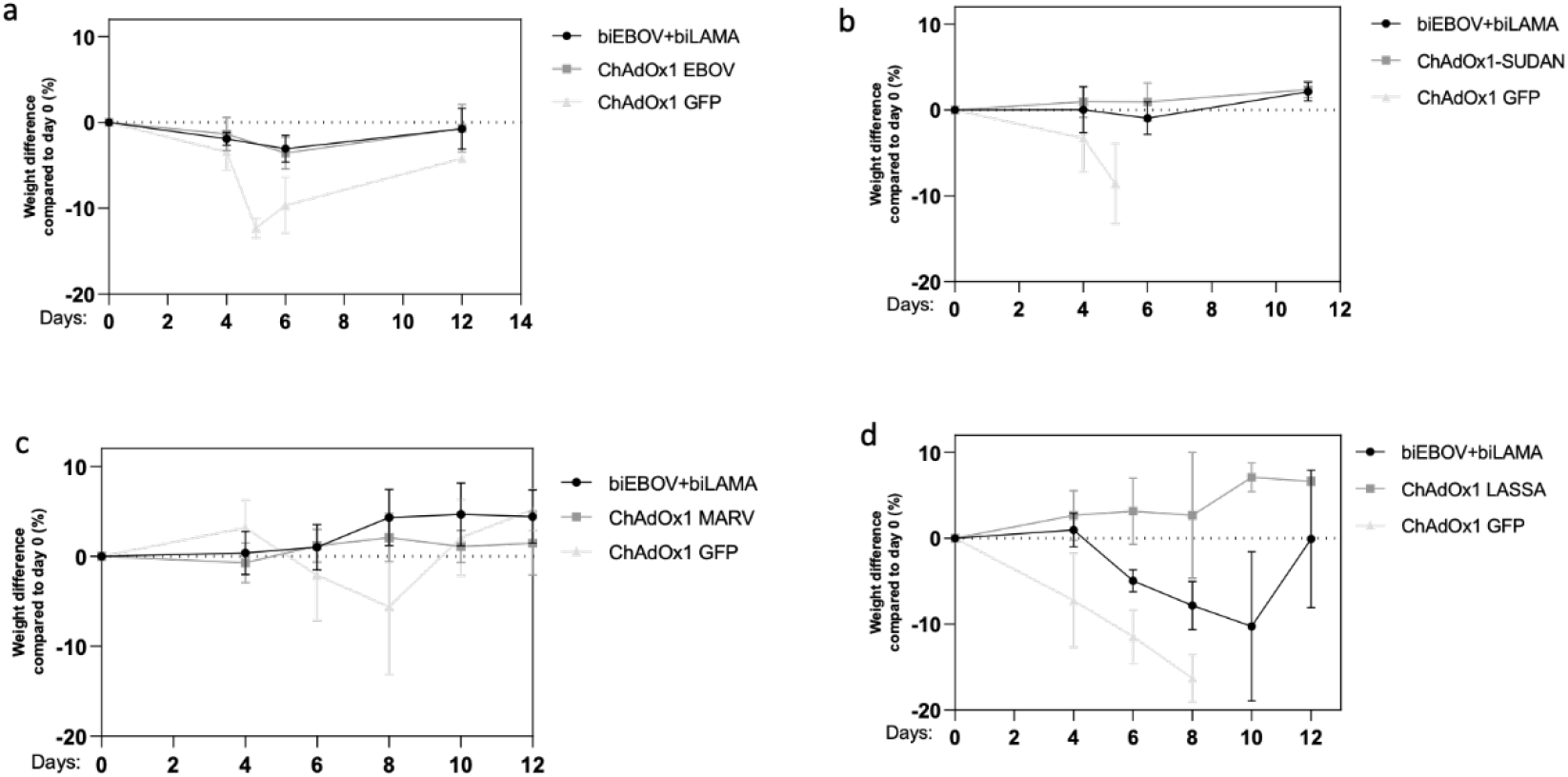
Weight loss after Ebola Zaire, Ebola Sudan, Marburg and Lassa virus infections in vaccinated mice. Groups of 6 IFNγ-/-mice received either biEBOV and biLAMA, relevant monovalent control or irrelevant control and were challenged 3 weeks later IP with virus. Animals were culled upon reaching humane end point. Weight loss for each of the challenge viruses: (a) Ebola Zaire Guinea Kissidoguo, (b) Ebola Sudan Boniface (c) Marburg Mukose and (d) Lassa Josiah. Mean and SD shown.

**Suppl Table 1.** Cellular immunogenicity in fold change of monovalent and ChAdOx1 bivalent vaccines. BALB/c mice received either bivalent vaccines or mix of monovalent controls and IFNγ ELISpot was performed at 2 weeks post vaccination. Vaccine regimens shown are: 1.biEBOV versus monovalent mix (EBOV & SUDV) 2. biLAMA versus monovalent mix (MARV & LASV) 3. biEBOV and biLAMA mixed versus monovalent mix (EBOV, SUDV, MARV, LASV) and 4. EBOV and biLAMA co-administered versus monovalent mix (EBOV, SUDV, MARV, LASV). Red indicates lowest fold change observed for regimen and green indicates highest.

| Vaccine regimens | Antigen | Fold change of Bivalent versus Monovalent regimens |
| --- | --- | --- |
| 1. biEBOV versus monovalent mix | EBOV GP | 0.88 |
|  | SUDV GP | 0.95 |
| 2. biLAMA versus monovalent mix | MARV GP | 0.93 |
|  | LASV GP | 1.14 |
| 3. biEBOV and BiLAMA mixed versus monovalent mix | EBOV GP | 0.36 |
|  | SUDV GP | 0.69 |
|  | MARV GP | 0.90 |
|  | LASV GP | 0.99 |
| 4. biEBOV AND BiLAMA co-administered versus monovalent mix | EBOV GP | 2.56 |
|  | SUDV GP | 1.53 |
|  | MARV GP | 1.27 |
|  | LASV GP | 1.31 |

**Suppl Table 2.**
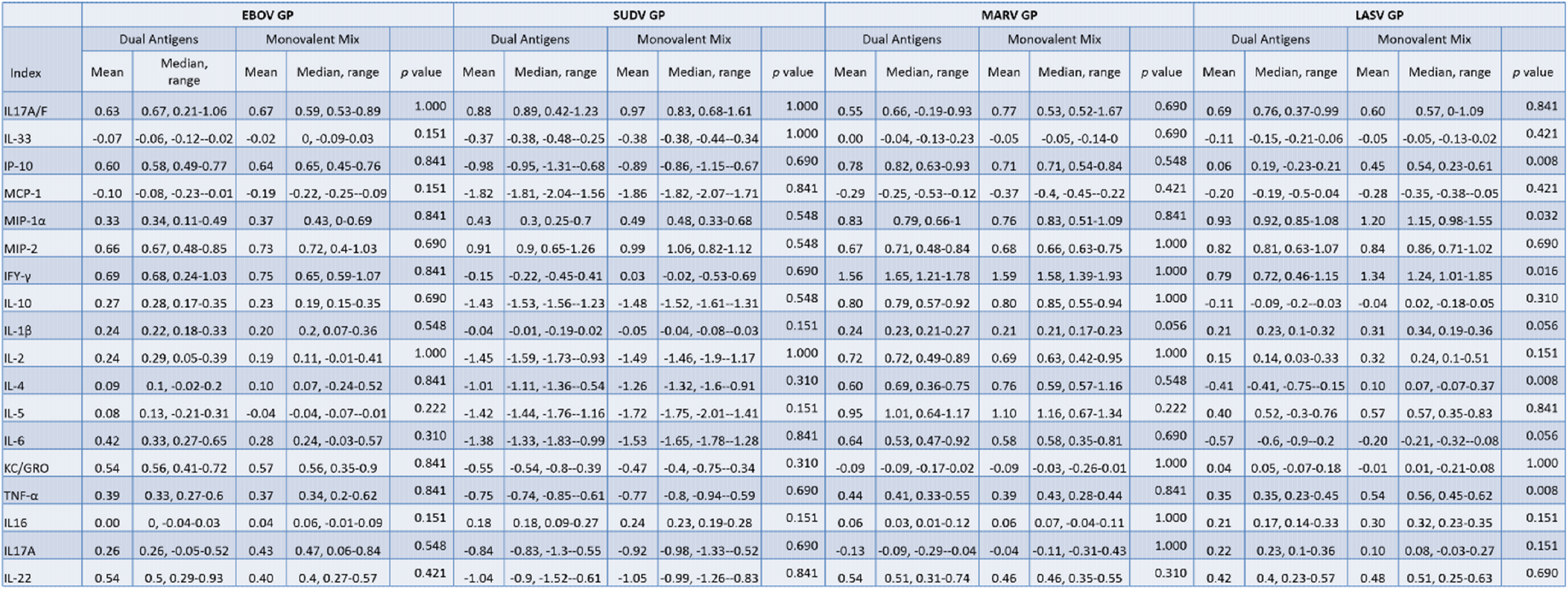
Cytokine and chemokine levels after prime-boost vaccination. BALB/C Mice were primed with either biEBOV and biLAMA co-administered or a mix of monovalent controls, 4 weeks later all mice were boosted with tetraMVA. Cytokine levels (in supernatant from splenocytes stimulated with the relevant peptide pools) determined using an MSD assay 2 weeks post boost. Log_10_ fold change in cytokine/chemokine levels in unstimulated versus peptide stimulated was compared across the vaccine groups. Values given to 2 d.p. for descriptives and 3 d.p. for p values (determined by Mann-Whitney)

